# Alteration of ribosome function upon 5-fluorouracil treatment favours cancer cell drug-tolerance

**DOI:** 10.1101/2020.06.04.131201

**Authors:** Gabriel Therizols, Zeina Bash-Imam, Baptiste Panthu, Christelle Machon, Anne Vincent, Sophie Nait-Slimane, Maxime Garcia, Mounira Chalabi-Dchar, Florian Lafôrets, Virginie Marcel, Jihane Boubaker-Vitre, Guillaume Souahlia, Marie-Alexandra Albaret, Hichem C. Mertani, Michel Prudhomme, Martin Bertrand, Jean-Christophe Saurin, Philippe Bouvet, Théophile Ohlmann, Jérôme Guitton, Nicole Dalla-Venezia, Julie Pannequin, Frédéric Catez, Jean-Jacques Diaz

**Author notes:** co-senior authors (Frédéric Catez, Jean-Jacques Diaz).

## Abstract

Partial response to chemotherapy leads to disease resurgence. Upon treatment, a subpopulation of cancer cells, called drug-tolerant persistent cells, display a transitory drug tolerance that lead to treatment resistance ^1,2^. Though drug-tolerance mechanisms remain poorly known, they have been linked to non-genomic processes, including epigenetics, stemness and dormancy ^2–4^. 5-fluorouracil (5-FU), the most widely used chemotherapy in cancer treatment, is associated with resistance. While prescribed as an inhibitor of DNA replication, 5-FU alters all RNA pathways ^5–9^. Here, we show that 5-FU treatment leads to the unexpected production of fluorinated ribosomes, exhibiting altered mRNA translation. 5-FU is incorporated into ribosomal RNAs of mature ribosomes in cancer cell lines, colorectal xenografts and human tumours. Fluorinated ribosomes appear to be functional, yet, they display a selective translational activity towards mRNAs according to the nature of their 5’-untranslated region. As a result, we found that sustained translation of *IGF-1R* mRNA, which codes for one of the most potent cell survival effectors, promoted the survival of 5-FU-treated colorectal cancer cells. Altogether, our results demonstrate that “man-made” fluorinated ribosomes favour the drug-tolerant cellular phenotype by promoting translation of survival genes. This could be exploited for developing novel combined therapies. By unraveling translation regulation as a novel gene expression mechanism helping cells to survive a drug-challenge, our study extends the spectrum of molecular mechanisms driving drug-tolerance.

## Main text

Translation regulation plays a major role in controlling gene expression and contributes to diseases emergence including cancer ^10,11^. Within ribosomes, ribosomal RNAs (rRNAs) play a central role in the translation process, by monitoring codon:anti-codon recognition, coordinating ribosomal subunit activity and catalysing peptide-bond formation through its ribozyme activity. rRNAs contain over 200 naturally occurring chemical modifications which stabilise rRNA structure and create additional molecular interactions not provided by non-modified nucleotides^12–14^. Chemical modifications of rRNAs were shown to directly contribute to translational regulation ^11,15,16^. We, and others, showed that rRNA chemical modifications contribute to the fine-tuning of ribosome functions and to modulating translational activity of ribosomes in cancer cells ^17–20^. 5-FU treatment results in 5-fluorouridine (5-Urd) incorporation into various types of cellular RNA including the precursor of rRNA ^9^. However, the consequences of 5-FUrd incorporation into ribosomal RNA precursor on ribosome production and functioning have so far not been analysed, neither is its impact on cellular phenotype.

### 5-FU does not inhibit ribosome production

Previous work indicated that at a high concentration, 5-FU alters ribosome biogenesis without inhibiting pre-rRNA synthesis ^8,21^. To further investigate this, we treated colorectal cancer HCT116 cells with clinically relevant concentrations of 5-FU (10-50 μM) ^22,23^, which result in growth inhibition and cell death (^24^ and Extended data Fig.1a). Within this concentration range, 5-FU treatment resulted in enlarged nucleoli, absence of nucleolar cap formation and absence of dispersion of nucleolar markers, as opposed to cells treated with the RNA Pol I inhibitor actinomycin D (Fig. 1a and Extended data Fig. 1b-c). Such nucleolar restructuring reveals an alteration of ribosome biogenesis albeit without pre-rRNA synthesis inhibition, and was confirmed by TEM (Extended data Fig. 1d). Consistently, 47S/45S pre-rRNA levels, analysed by Northern blotting and RNA fluorescent *in situ* hybridization (FISH), were unchanged following 5-FU treatment confirming that 5-FU did not affect RNA Pol I activity (Fig. 1b and Extended data Fig. 1e and Fig. 2a-b).

**Fig. 1.**
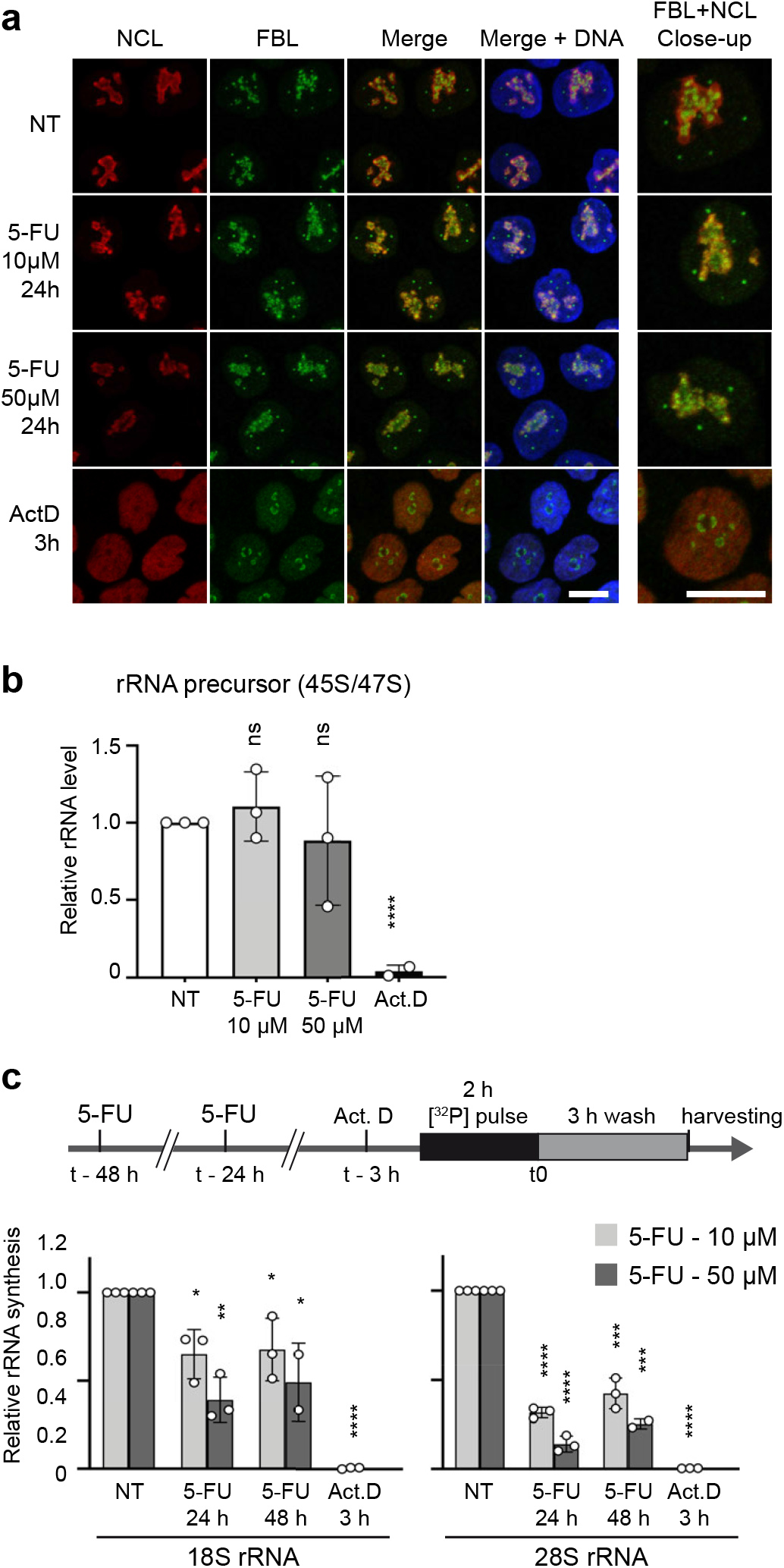
Ribosome production is maintained in 5-FU-treated cells. HCT116 cells were treated with 5-FU at 10 μM or 50 μM for 24 h or 48 h or with actinomycin D (Act.D) for 3 h as a reference of rRNA synthesis inhibition. **a**, Morphology of nucleoli analysed by immunofluorescent detection of nucleolar markers nucleolin (NCL, red) and fibrillarin (FBL, green). Nuclei were stained with Hoechst (blue) Scale bar = 10 μM. **b**, Pre-rRNA synthesis analysed by detection of 47S/45S rRNA precursor levels by Northern blotting. Data are expressed as mean values +/- s.d. of independent experiments (n = 3). **c**, Rate of 28S and 18S rRNAs production analysed by isotope pulse labelling. Radioactivity was measured for each rRNA and normalised against ethidium bromide. Data are expressed as mean +/- s.d. of independent experiments (n = 3). Results of unpaired two-tailed t-test are indicated as non-significant (ns) p<0.05 (*), p<0.01 (**), p<0.001 (***) and p<0.0001 (****).

Northern blot analysis also confirmed that ribosome maturation at post-transcriptional steps was altered, and revealed that the pre-rRNA processing was impaired at the cleavage stage at site 2 (Extended data Fig. 2a-c). Yet, despite this effect, the late pre-rRNA intermediates leading to 18S and 28S rRNA were still detected (Extended data Fig. 2c) suggesting that ribosome production was in part maintained. This was confirmed by [^32^P] pulse-chase experiments that showed that ribosomes are produced at significant levels for up to 48 h under 5-FU treatment (Fig. 1c and extended data Fig. 2d). Thus, at clinically relevant concentration of 5-FU, each step of ribosome processing is able to proceed, despite the stringent quality control, thus allowing ribosome production to be maintained at a substantial level.

### 5-FU incorporation into ribosomes

5-FU was previously shown to be incorporated in RNAs ^9^. We therefore wondered whether ribosomes produced and exported to the cytoplasm in treated cells contain 5-FUrd within their rRNA. To this end, we developed a quantitative LC-HRMS approach that now allows us to determine the number of 5-FU incorporated in rRNA of cytoplasmic ribosomes purified at high stringency (Fig. 2a, see methods for details ^25^). We found that ribosomes contained significant amounts of 5-FUrd, ranging from 7 to 15 5-FUrd molecules per ribosome upon 24 h of treatment with 5 μM to 100 μM of 5-FU (Fig. 2b). We ruled-out that the 5-FU signal came from non-ribosomal RNA by measuring 5-FUrd from gel-purified 18S and 28S rRNA (Extended data Fig. 3a). 5-FU was incorporated into rRNA from cytoplasmic ribosomes purified from a panel of cell lines representing cancers for which 5-FU-based therapies are commonly used, including colorectal, pancreatic and triple negative breast cancers (Fig. 2c). Altogether, these data demonstrate that upon 5-FU treatment, ribosomes containing fluorinated rRNA are fully assembled and exported to the cytoplasm, showing that presence of 5-FUrd is tolerated by the quality control systems of the cell.

**Fig. 2.**
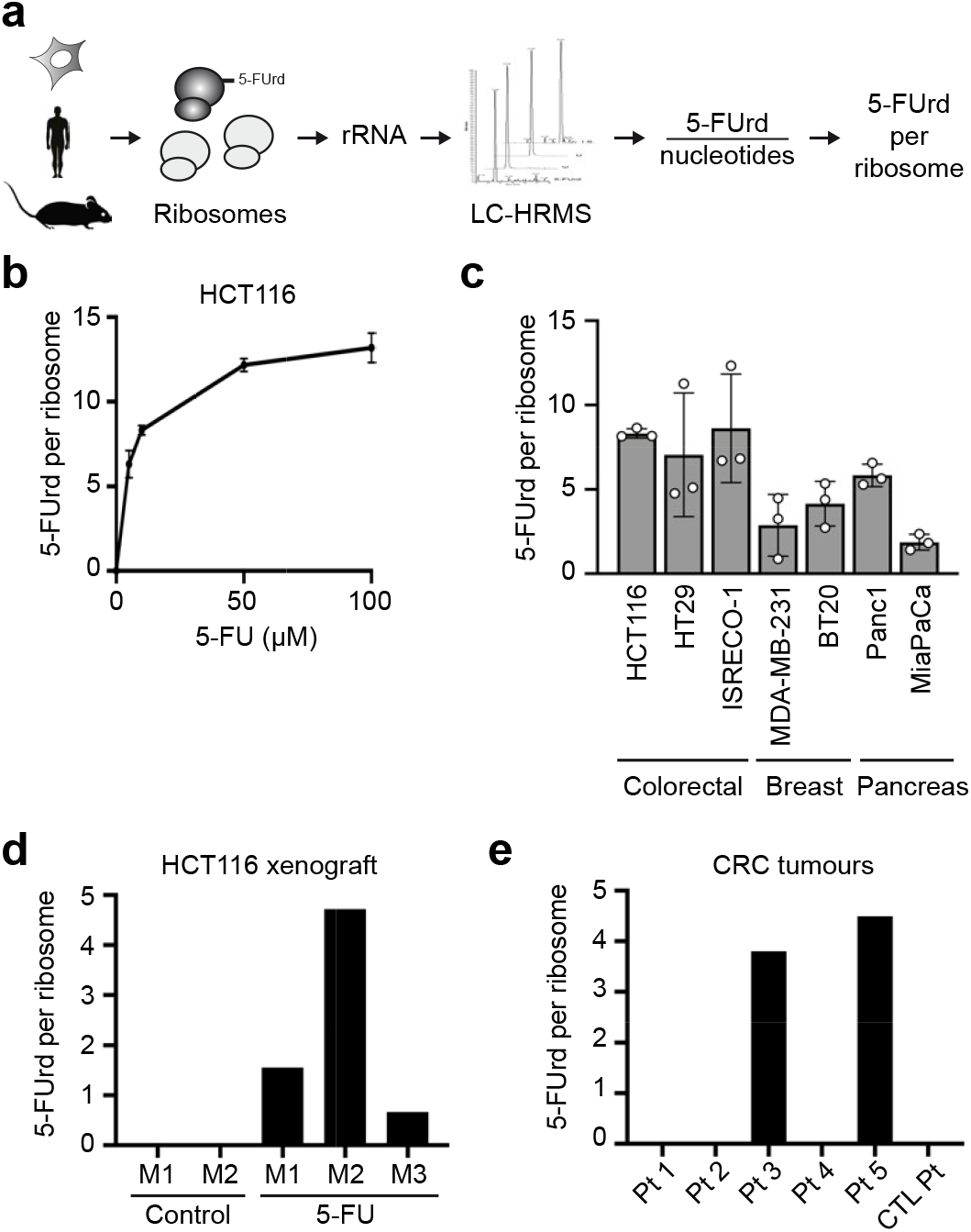
5-FU is incorporated in ribosomes of cell lines and tumours. **a**, Schematic representation of 5-FUrd incorporation into ribosomes determined using LC-HRMS. **b**, HCT116 cells were treated for 24 h with 5 to 100 μM 5-FU and 5-FUrd incorporation was determined as in **a**. Data are expressed as mean +/- s.d. of independent experiments (n = 3). **c**, Indicated cell lines were treated for 24 h with 10 μM of 5-FU and incorporation of 5-FUrd into rRNA was determined as in **a**. Data are expressed as mean +/- s.d. of independent experiments (n = 3). **d**, HCT116 cells were xenografted into nude mice, and mice were treated with 50 mg/kg of 5-FU twice a week (5-FU) or with PBS (Control). Incorporation of 5-FUrd into rRNA was determined as in **a**. Data are values for individual animals (n = 1). **e**, rRNA were purified from total RNA extracted from colorectal cancer samples. Incorporation of 5-FUrd into rRNA was determined as in **a**. Pt = sample from 5-FU treated patient, CT Pt = sample from patient not treated with 5-FU. n = 1 for each sample.

Next, we investigated whether fluorinated ribosomes could be produced within tumours *in vivo*. First, we analysed rRNA from HCT116 xenografts established in nude mice. 5-FU treatment efficacy was evidenced by a decrease in tumour growth (Extended data Fig. 3b). 5-FU was detected in mature rRNA purified from tumour cells collected after the last treatment at levels close to those observed in cultured cells (Fig. 2d). Thus, 5-FU incorporation in ribosomes can be replicated in a common xenografted animal model. Finally, we analysed rRNA of colorectal tumour cells from patients treated with 5-FU-based therapies, using large RNA quantities to optimise detection (Extended data Fig. 3c). Of the 5 samples tested from 5-FU-treated patients, 5-FUrd was detected in rRNA from 2 patients (3.80 and 4.50 5-FUrd per ribosome respectively; Fig. 2e), a patient receiving no 5-FU serving as a negative control. Altogether, these data show that 5-FUrd incorporates into rRNA of cells treated with 5-FU, and that 5-FU-based chemotherapy leads to the production of fluorinated ribosomes within tumour cells in animal models and human.

### Altered translation by Fluorinated ribosomes

Because rRNAs and their post-transcriptional chemical modifications play a central role in ribosome functioning, and because 5-FU induces changes in translational regulation ^24,26,27^, we postulated that fluorinated ribosomes may display modified translational activity. To investigate this, we first considered whether fluorinated ribosomes could be recruited onto mRNA during translation, by analysing the rRNA 5-FU content in actively translating ribosomes isolated by sucrose gradient. 5-FU was readily detected in actively translating ribosomes, demonstrating that fluorinated ribosomes can engage in translation (Fig. 3a). Next, we evaluated whether incorporation of 5-FU in rRNA impacts the translational capacity of ribosomes. We used our recently developed *in vitro* hybrid translation assay ^19,28^, in which ribosomes alone have been exposed to 5-FU, in order to evaluate the activity of purified fluorinated ribosomes in a controlled setting (Fig. 3b). The translational capacity of fluorinated ribosomes was assessed using a set of luciferase reporter mRNAs, the translation of which relies on different 5’UTR: (i) short 5’UTR from globin and GAPDH mRNAs, (ii) long and structured capped 5’UTR from IGF-1R and c-Myc mRNAs, and (iii) long and structured uncapped 5’UTR from viral mRNA of cricket paralysis virus (CrPV) and encephalomyocarditis virus (EMCV), which initiate translation through an internal ribosome entry site (IRES). The results showed first that fluorinated ribosomes were not impaired for translation, and second that they displayed a selective translation initiation efficacy that differed from that of control ribosomes, and varied according to the nature of the 5’UTR of the reporter mRNA used (Fig. 3c). Indeed, first, globin and GAPDH were less efficiently translated, a result that is consistent with lower overall protein synthesis in 5-FU treated cells (^24^, and Extended data Fig. 4a). Second, reporter mRNAs containing IGF-1R and c-Myc 5’-UTR were more efficiently translated by fluorinated ribosomes. These differences suggest that translation efficiency varies according to the nature of the 5’UTR indicating that the initiation step of translation was different for fluorinated ribosomes compared to normal ribosomes. To consolidate this hypothesis, translation was tested on a mRNA carrying the CrPV intergenic IRES, an element that directly binds to the ribosome and initiates translation without any cellular translation initiation factors (eIFs). Fluorinated ribosomes displayed a decrease translational activity on CrPV mRNA, strongly supporting that fluorinated ribosomes are structurally or functionally different (Fig. 3d). This defect in translation initiation from the CrPV intergenic IRES was not strictly related to cap-independent initiation mechanisms since fluorinated ribosomes were more efficient at translating an EMCV IRES containing mRNA, another cap-independent translation initiation model (Fig. 3d). Altogether, these experiments demonstrate that 5-FU incorporation into rRNA modifies the ability of ribosomes to initiate mRNA translation from different 5’UTR, and highlight that fluorinated ribosomes might contribute to 5-FU induced translational reprograming that we previously observed ^24^.

**Fig. 3.**
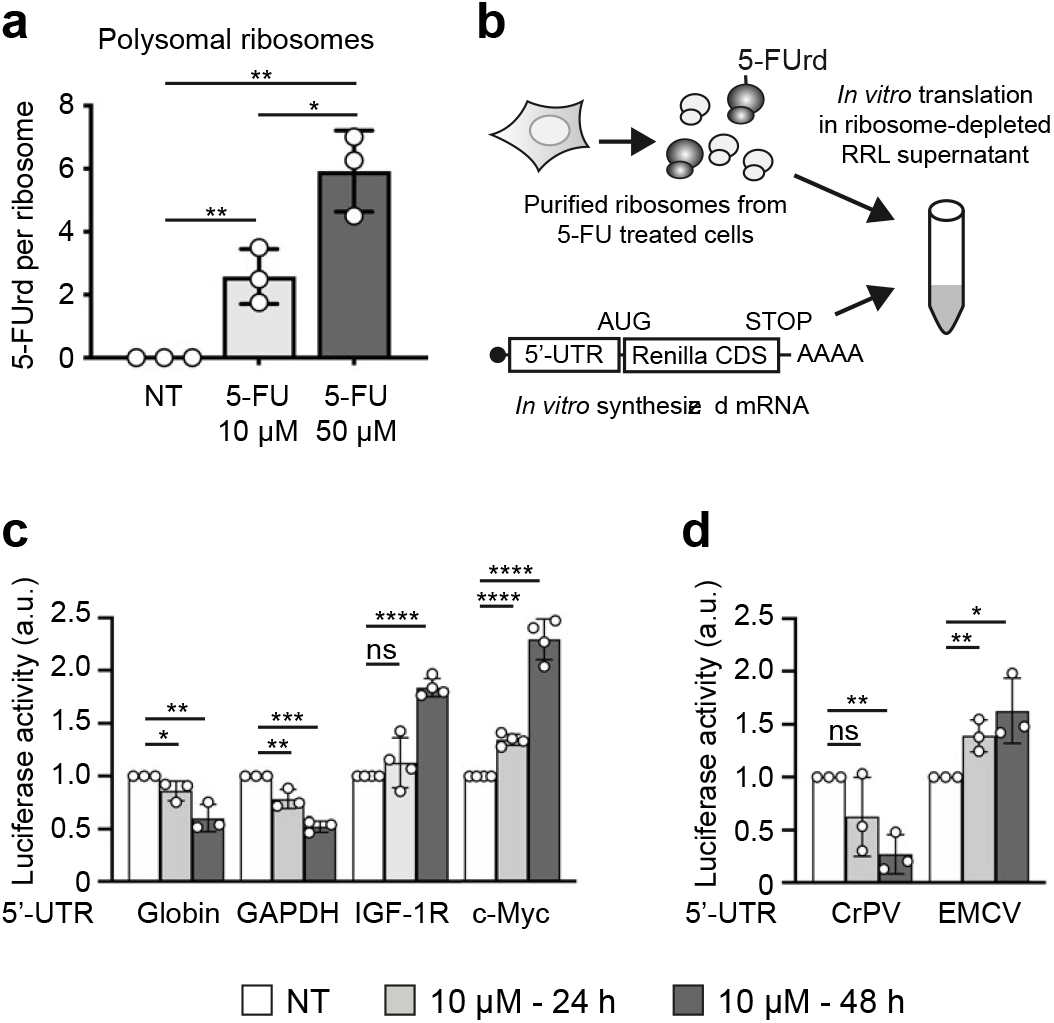
Fluorinated ribosomes display altered translational properties. **a**, HCT116 cells were treated for 24 h with either 10 μM or 50 μM 5-FU, and translationally active ribosomes were purified from the polysomal fraction. Incorporation of 5-FUrd was measured by LC-HRMS. Data are expressed as mean +/- s.d of independent experiments (n = 3). **b**, Schematic representation of the hybrid *in vitro* translation assay used in **c** and **d**. **c, and d**, Ribosomes were purified from HCT116 cells treated with 10 μM 5-FU for 24 h or 48 h, and their translational activity was evaluated using the hybrid *in vitro* translation assay. Translation efficacy was evaluated on luciferase reporter mRNA containing the 5’-UTR of the indicated gene. Values are units of Renilla luciferase activity normalised against the untreated (NT) condition. **c**, Evaluation on capped mRNA containing the 5’UTR of actin, GAPDH, IGF-1R and c-Myc genes. **d**, Evaluation on uncapped mRNA containing the IRES element from the cricket paralysis virus (CrPV) and the encephalomyocarditis virus (EMCV). Data are expressed as mean +/- s.d. of independent experiments (n = 3). Results of unpaired two-tailed t-test are indicated as non-significant (ns) p<0.05 (*), p<0.01 (**), p<0.001 (***) and p<0.0001 (****).

### IGF-1R promotes 5-FU drug-tolerance

The data above suggest that fluorinated ribosomes favour translation of selected mRNAs, including genes such as *IGF-1R* and *c-Myc*, that may promote early cell survival and lead to resistance ^4,29,30^. We focused on *IGF-1R*, a gene that play a major role in tumorigenesis and whose contribution to cell survival has been largely demonstrated in various models including colorectal cancer ^31–34^. Because 5-FU treatment induces a decrease in global protein synthesis (^24^, Extended data Fig. 4a), we initially evaluated whether IGF-1R mRNA translation was also impacted by 5-FU treatment in HCT116 cells. *IGF-1R* mRNA translation efficacy was assessed by measuring the recruitment of cytoplasmic mRNA into the heavy polysome fraction of control and 5-FU-treated cells (Fig. 4a). Our data show that the fraction of *IGF-1R* mRNAs associated with heavy polysomes was maintained in 5-FU treated cells, while that of actin and GAPDH mRNAs decreased, indicating that translation of *IGF-1R* mRNA remains largely unaltered upon 5-FU treatment. Consistently, the global level of IGF-1R protein was also maintained in treated cells (Fig. 4b and Extended data Fig. 4b). Next, to determine whether the IGF-1/IGF-1R pathway contributes to the survival of CRC cells exposed to 5-FU, cells were first treated with 5-FU for 24 h or 48 h, and were subsequently treated with IGF-1. Cell proliferation was monitored over 5 days, and revealed that while IGF-1 had no impact on control cells, it improved the growth of cells treated with 5-FU (Fig. 4c and d). To further validate our findings, HCT116 cells were co-treated with 5-FU and the IGF-1R kinase inhibitor NVP-AEW541 ^35^, and cell response was monitored by cell counting using high-content analysis (Fig. 4e). Inhibition of IGF-1R further decreased the number of cells that survived 5-FU treatment, an observation that was confirmed by MTS assay (Extended data Fig. 4d), demonstrating that an active IGF-1/IGF-1R pathway is necessary for optimal cell tolerance to 5-FU. Overall, our results unveil that the IGF-1/IGF-1R pathway plays a role in the survival of a cell subpopulation upon 5-FU treatment, and strongly support that the 5-FU driven maintenance of IGF-1R synthesis contributes to this mechanism.

**Fig. 4.**
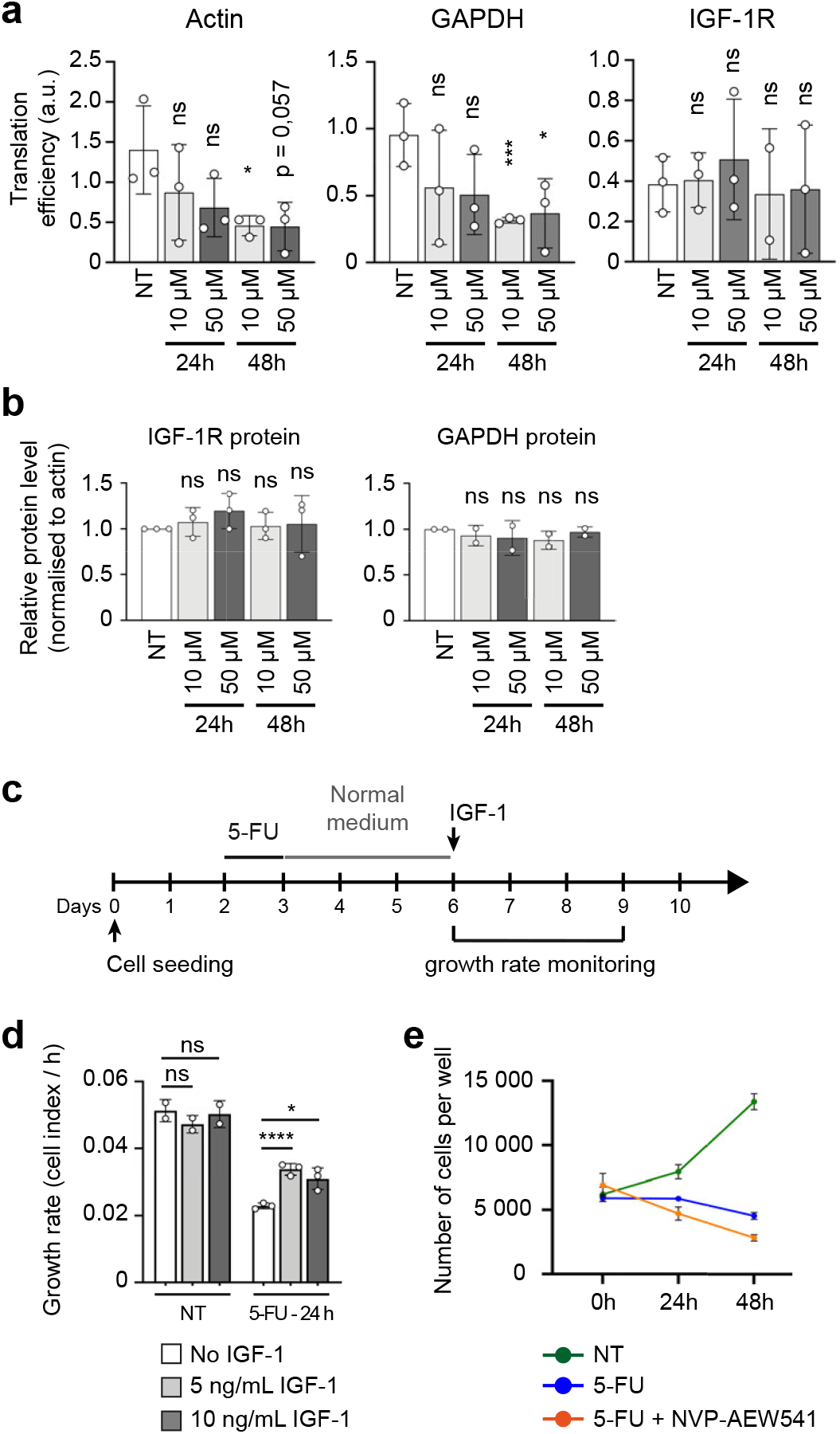
IGF-1R contributes to survival and recovery of 5-FU treated CRC cells. **a, b**, HCT116 cells were treated with 10 μM or 50 μM 5-FU for 24 h or 48 h or untreated (NT). **a**, Translation efficiency of actin, GAPGH and IGF-1R mRNAs. Each mRNA was quantified from cytoplasmic and polysomal fractions. Translation efficiency are shown as the ratio of polysomic mRNA over the cytoplasmic mRNA. Each dot represents an individual biological sample measured in duplicate and data are expressed as mean ± s.d of independent experiments (n = 3). **b**, Level of IGF-1R protein (left) and GAPDH protein (right) normalised against the Ku80 housekeeping gene, quantified from Western blot. Each dot represents an individual biological sample and data are expressed as mean ± s.d of independent experiments (n = 3). **c, d**, HCT116 cells were treated with 10 μM 5-FU for 24 h or 48 h or NT, and not stimulated (No IGF-1) or stimulated with 5 or 10 ng/mL of IGF-1. Cell growth was monitored in real-time over 5 days. **c**, Schematic representation of the experiment. **d**, Growth rate measured over 72 h (day 6 to day 9). Each dot represents a technical replicate and data are expressed as mean ± s.d. **e**, HCT116 cells were treated with 10 μM of 5-FU alone or with 5 μM of IGF-1R inhibitor NVP-AEW541 for 48 h, or NT. The number of cells per well was counted by image analysis. Each dot represents a technical replicate and data are expressed as mean ± s.d. of independent experiments (n = 3). Results of unpaired two-tailed t-test are indicated as non-significant (ns) p<0.05 (*), p<0.01 (**), p<0.001 (***) and p<0.0001 (****).

## Discussion

In this study, we reveal that the pyrimidine analogue 5-FU is incorporated into ribosomes *in vitro* and *in vivo*, including in human tumours. We used a novel LC-HRMS method that we developed ^25^ in order to quantitate the level of incorporation of 5-FUrd in a defined RNA molecule. This approach allowed us to demonstrate that 5-FUrd is incorporated into ribosome at significant levels, showing that cells can tolerate the production of non-natural ribosomes. This finding was unexpected because ribosome assembly and maturation are under stringent quality-control that induces the degradation of unproperly folded and assembled rRNAs ^36^, as evidenced by the decrease in the level of the late pre-rRNA species that we report in this study. As a result, cytoplasmic functional ribosomes contained up to 15 copies of 5-FUrd per ribosome, a number likely underestimated since only a fraction of the ribosome population was renewed within the time frame of our experiment. While addition of fluorine into rRNA results is a non-natural modification, and could be anticipated as deleterious, we found that fluorinated ribosomes are functional as they engage in translation. However, their activity is altered and displays a selective ability to initiate mRNA translation according to the nature of its 5’UTR. Hence, fluorine appears to modify the functioning of the rRNA, since (i) chemical modifications of rRNA including 2’-O-methylation and pseudouridylation were shown to contribute to translational regulation and efficiency ^18,37–39^, and (ii) structural studies of bacterial and human ribosomes showed that chemically-modified nucleotides establish molecular interactions that cannot be provided by non-modified ones ^12,14^. In particular, the presence of 5-FUrd in the functional domains of the ribosome, such as the A, P and E sites are more likely to impact translation. The fine mapping of the location of 5-FUrd within rRNA may improve our understanding of its impact on ribosome functioning at the atomic level.

We previously described a major translational reprograming induced by 5-FU in colorectal cancer cells, that we have linked to a miRNA-based mechanism ^24^. The fluorination of rRNA that we describe herein represents an additional mechanism by which 5-FU contributes to translational reprograming of treated cells ^24^. It is likely that other mechanisms are involved, such as 5-FUrd incorporation into mRNAs.

We determined that the 5-FU altered translational machinery contributes to maintaining the expression level of the IGF-1R gene, thus promoting cell survival. This suggests that cytotoxic efficiency of 5-FU may be improved if fluorinated ribosome production is prevented, an approach that could be effectively tested using the recently developed ribosome biogenesis inhibitors, for which anti-cancer activities are being unveiled ^40–42^. These inhibitors have been positively evaluated in the context of Myc-driven pathologies, lymphoma and in combination with mTOR inhibitors. Our data now indicate that this novel class of drugs may be useful for improving non-targeted therapies.

Drug-tolerance is a critical phase as it represents a window of opportunity for genetic and non-genetic events to take place and provide cells with a drug-resistant phenotype. We show that sustained IGF-1R synthesis is a significant factor for cell survival upon 5-FU treatment. Surprisingly, our data indicate that 5-FU sensitized cells to IGF-1. It is not clear whether this is directly related to changes in translational regulation, nevertheless, it suggests that targeting the IGF-1/IGF-1R pathway may improve 5-FU efficacy. To our knowledge, this is the first line of evidence that the IGF-1/IGF-1R pathway might contribute to drug-tolerance. 5-FU is the most widely used chemotherapy, and there is a high demand for improving its efficacy. Our data highlight the potential benefits of understanding drug-tolerance mechanisms in response to 5-FU, which has so far not been fully described. In addition, while our study focused on a base analogue incorporated into RNA, other compounds binding to RNAs such as platin derivatives or any drug that might interfere with RNA metabolism should now also be considered as modifiers of ribosome structure and activity ^43^, and may also contribute to altering translational regulation in treated cells, with a deleterious impact for patient outcome.

Altogether, our study extends the spectrum of gene expression mechanisms that help cells survive a drug-challenge, by adding translational regulation to epigenetics, stress response, metabolism adaptation and stemness or dormancy phenotypes ^1,2,4,44–46^. These findings also reveal that exposure to drugs can result in the production of new “man-made” biological complexes, the functioning of which cannot be anticipated, and that require further studies to fully comprehend drug response and propose new therapeutic strategies.

## Acknowledgements

We thank E. Errazuriz (Centre d’Imagerie Quantitative Lyon-Est) and C. Vanbelle (plateau d’imagerie du CRCL) for technical help. We thank Brigitte Manship for editing the manuscript. G.T. and S.N-S. were recipient of a Ph.D. fellowship from la Ligue Nationale Contre le Cancer and from “La Fondation pour la Recherche Médicale”. MG was supported by ANR Grant “INVADE” PC201507. The project was supported by the “Ligue Contre le Cancer, comités départementaux de la Drôme, du Rhône, du Puy-de-Dôme, et de l’Allier”, by Agence Nationale de la Recherche (grant RiboMeth - ANR-13-BSV8-0012), by the Institut National du Cancer (INCa, grant FluoRib - PLBIO18-131) and by Programmes d’Actions Intégrées de Recherche (PAIR Sein, RiboTEM, ARC_INCa_LNCC_7625). NDV, AV, JP and FC are CNRS research fellow. JJD, and VM are Inserm research fellows.

## Author contributions

G.T., Z. B-I., B.P., J.G., C.M., A.V., S.N-S., M.G., M. C-D., F.L., V.M., G.S., S.H., M-A. A., J.B.V., J.P. and F.C. performed experiments. G.T., J.G., V.M., H.C.M., J.C.S., N.D.V., T.O., J.P., F.C. and J-J.D. designed experiments. M.P. and M.B. provided human biopsies. T.O., J.P., J.G., N.D.V., F.C and J-J.D. designed and supervised the study. F.C. and J-J.D. wrote the paper.

## Data availability

All data and materials are available from the authors upon reasonable request.

---

**Extended data Table 1:**
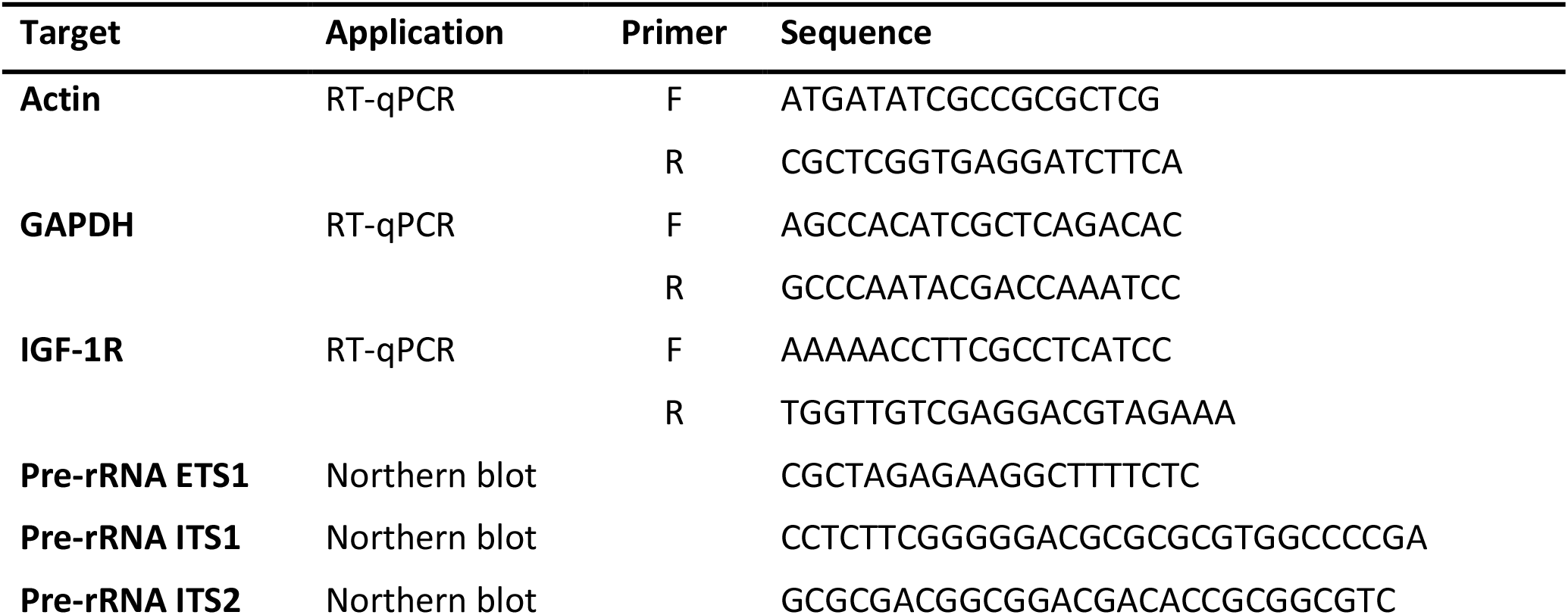
Sequence of oligonucleotides used in this study

