## Supplementary data and methods for "Alteration of ribosome function upon 5-fluorouracil treatment favours cancer cell drug-tolerance"

**Altered translation by fluorinated ribosomes favours 5-FU drug-tolerance**

*co-senior authors

### SUPPLEMENTARY MATERIAL

### METHODS

#### Cell lines, Cell culture and 5-FU treatment

Cells were maintained in Dulbecco Minimum Essential Medium – GlutaMax (Invitrogen) supplemented with 10% foetal bovine serum (FBS) at 37°C with 5% $CO_2$. The following cell lines were obtained from ATCC: HCT116 (ATCC CCL-247), MDA-MB-231 (ATCC HTB-26), BT20 (ATCC HTB-19), HT29 (ATCC HTB-38), Panc1 (ATCC CRL-1469) and MiaPaCa (ATCC CRL-1420). The following cell lines were obtained from the authors: ISRECO1 (Cajot, J. F., et al. Cancer Res. (1997) 57, 2593–2597). The following cell lines were authenticated by 21 PCR-single-locus-technology (Eurofins, Ebersberg, Germany): HCT116, HT29, MDA-MB-231, MiaPaCa, Panc1. ISRECO1 cell line was not authenticated by PCR-single-locus-technology as this cell line genetic pattern is not described in databases. Cells were routinely tested against mycoplasma infection.

Cells were plated 48h before 5-FU treatment. 5-FU was kindly provided by the Centre Léon Bérard (Lyon, FRANCE) and was purchased from Sanofi-Aventis. The stock solution was diluted immediately before use in DMEM.

#### Western Blot

Western blot was performed as previously reported [1]. Briefly, cells were lysed in lysis buffer A (20 mM HEPES-KOH pH 7.2, 100 mM KCl, 1 mM DTT, 0.5 mM EDTA, 0.5 % NP40, 10 % Glycerol) supplemented with protease inhibitor (Complete EDTA free, Roche). Proteins were subjected to SDS-PAGE, and blotted onto poly-vinylidene difluoride (PVDF) membranes (Immobilon-P, Millipore). Membranes were blocked in Tris-buffered saline solution containing 0.05 % Tween 20 and 5 % non-fat milk and incubated with primary antibodies: mouse monoclonal antibodies against Ku80 (ab119935, Abcam), GAPDH (AM4300, Invitrogen) and rabbit monoclonal antibodies against IGF-1R (#9750, Cell Signalling) and actin (ab179467, Abcam). Horseradish peroxidase conjugated secondary antibodies (Cell Signalling) were used for the detection of immunoreactive proteins by chemiluminescence (Clarity Western ECL Substrate, BioRad)). Imaging and densitometric measurements of the bands were performed using Image lab software (BioRad).

**Ribosome purification from whole cells**

Ribosome purification was performed as previously described [2]. Briefly, cytoplasmic fractions were obtained by mechanical lysis of cells using a Precellys tissue homogenizer (Ozyme) and centrifugation at 12,000 g for 10 min to pellet mitochondria. To purify ribosomes, cytoplasmic fractions were loaded onto a 1 M sucrose cushion in a buffer containing 50 mM Tris-HCl pH 7.4, 5 mM $MgCl_2$, 500 mM KCl and 2 mM DTT, and centrifuged for 2 h at 240,000 g. The pellet containing the ribosomes was resuspended in a buffer containing 50 mM Tris-HCl pH 7.4, 5 mM $MgCl_2$ and 25 mM KCl.

**Ribosome purification from polysomal fraction**

Cells were seeded at $10^7$ cells/15 cm dish and treated with 5-FU 48 h later as indicated. Cells were then incubated for 5 min with 25 µg/mL Emetin (Sigma) and washed twice with cold 1X PBS before harvesting. Cytosolic lysates were prepared as described above. 3 mg of cytosolic proteins was loaded onto a 10-40 % sucrose gradient, sedimented by ultra-centrifugation for 2 h at 240,000 g at 4°C on a SW-41 rotor (Beckman). Fractions were collected and absorbance profiles were generated at 245 nm using an ISCO UA-6 detector. Polysomal fractions were pooled and concentrated in an Amicon Ultra-15 unit with a 100 kDa cut-off. KCl concentration was adjusted to 500 mM using a 4 M stock solution. The ribosome suspension was reduced to 600 µL on an Amicon Ultra-15 filtration unit, and loaded onto a 1 M sucrose cushion in a buffer containing 50 mM Tris-HCl pH 7.4, 5 mM $MgCl_2$, 500 mM KCl and 2 mM DTT, and centrifuged for 2 h at 240,000 g. The pellet containing the ribosomes was resuspended in a buffer containing 50 mM Tris-HCl pH 7.4, 5 mM $MgCl_2$ and 25 mM KCl.

**rRNA purification**

For purified ribosomes, RNA was extracted using the TriPure Reagent (Roche) according to the manufacturer's instruction. Purified rRNAs were resuspended in water and quantified by spectrophotometry. For xenograft and human tumour samples, rRNAs were purified as described previously [3]. Briefly, 10 µg total RNA was denatured in 50% formamide at 70°C for 10 min, and separated on a 0.8% low-melting agarose gel in 0.5X TAE buffer. 18S and 28S rRNA were gel-purified using the NucleoSpin Gel and PCR Clean-up kit and NT1 buffer (Macherey Nagel) according to the manufacturer's instruction.

**5-FU analysis by LC-HRMS**

Purified rRNA (1 to 3 µg) was digested overnight at 37°C with 270 units of Nuclease S1 (Promega) using the supplied buffer. Next, nucleotides were dephosphorylated by directly adding to the mix 5U of calf intestine phosphatase (New England Biolabs) in 100 mM Tris-HCl, 50 mM NaCl, 10 mM $MgCl_2$, 0.025% Triton® X-100. Digestion was carried out overnight at 37°C, the digested mix was then stored at -80°C. For LC-HRMS analysis, on the day of the analysis, 300 µL of a mixture of methanol / water (70/30; v/v) was added and the samples were vigorously vortexed following the addition of labelled internal standard. Samples were centrifuged for 5 min. at 13,000 g, the supernatants were separated and evaporated to dryness under nitrogen at 37°C. The residues were then resuspended in water before injection into the mass spectrometer device. Analysis was carried out on a liquid chromatography coupled with a high-resolution mass spectrometer (Q-Exactive Plus Orbitrap - Thermo Scientific). Data acquisition was performed alternatively in negative and positive modes with the full scan mode (FS) at a resolution of 70,000 or 140,000 and with the parallel reaction monitoring mode (PRM) at a resolution of 17,500. Chromatographic separation of nucleosides was achieved on a Hypercarb® column ((2.1 mm x 100 mm; 5 µm) (ThermoScientific)). The level of 5-FU per ribosome was calculated as the ratio of measured [5-FUrd] over the measured [A], [C] and [G], divided by the relative quantity of each nucleotide per ribosome.

**Immunofluorescence analysis**

Cells were grown on glass coverslips, fixed in 4% of paraformaldehyde in phosphate buffered saline (PBS) before permeabilization with 0.5% Triton X-100 in PBS. Fibrillarin, Dyskerin and Nucleolin were detected using the anti-FBL rabbit polyclonal antibody (ab5821, Abcam) diluted at 1:2,000, anti-DKC1 rabbit polyclonal (sc-48794, Santa Cruz Biotechnology) diluted at 1:500 and anti-NCL mouse monoclonal antibody (ab13541, Abcam) at 1:4,000. Secondary antibodies were labelled with AlexaFluor 488 or AlexaFluor 555 (Molecular Probes) and used at 1:1,000. Coverslips were mounted using the Fluoromount G mounting medium (EMS). Images were acquired on a Zeiss LSM 780 confocal microscope using a 63X Plan Apochromat immersion objective (NA 1.4), as a Z-stack (voxel size: 0.0634 x 0.0634 x 0.3155 µm). Final images were prepared by maximum intensity projection to display all nucleoli, using Zeiss ZEN Black software. As indicated, level images of actinomycin D treated cells was adjusted to correct for NCL dispersion. Images were cropped using ImageJ (https://imagej.net) [4].

**Electron microscopy**

Cells were grown in a 6-well plate for 24 h and fixed with 2% glutaraldehyde (Sigma) in 0.1 M Sodium Cacodylate buffer at room temperature for 30 min. After washing three times in 0.2 M Sodium Cacodylate buffer, cell cultures were post-fixed with 2% aqueous Osmium Tetroxide (Sigma) and dehydrated in a graded series of ethanol at room temperature and embedded in Epon (Polysciences). After polymerization, ultrathin sections were collected on 300 mesh grids coated with Formvar (SPI supply) and stained with aqueous 1% uranyl acetate (SPI supply) and citrate (Leïca Ultrostainer) before observation on a Philips CM 120 transmission electron microscope at an acceleration voltage of 80 kV.

**Northern blot analysis**

Northern blot was performed as described in [5]. The probes were obtained by oligonucleotide synthesis (Eurogenetec) and are described in Extended data Table 1. 50 pmoles of each oligonucleotide probe was labelled in the presence of 50 pmoles of [$\gamma$-$^{32}$P] ATP (Perkin Elmer) and T4 polynucleotide kinase (NEB) for 30 min at 37°C. 3 µg of nuclear RNAs were resolved on a 1% denaturing agarose gel and blotted onto a Hybond-N+ membrane (GE Healthcare). Signal detection was performed using a PhosphorImager (FLA 9500, GE Healthcare). Total 28S and 18S rRNA were visualised by fluorescence imaging following ethidium bromide staining and were used as loading controls. Radioactivity was measured for each band and normalised against 18S and 28S rRNA signals stained with ethidium bromide as loading references. Quantification was performed using ImageJ (https://imagej.net) [4].

**[$^{32}$P] pulse-chase labelling**

Labelling of newly synthesised RNAs was performed as described previously [6]. Briefly, cells were grown in DMEM containing 10% FBS, and 5-FU containing medium for 24 h or 48 h or actinomycin D at 50 ng/mL for 3 h. Phosphate deprivation was performed by incubating cells for 1 h with phosphate-free DMEM (Invitrogen) containing 10% dialysed FBS. For labelling, cells were incubated with phosphate free-DMEM containing 10% dialysed FBS and 14 µCi/mL [$^{32}$P]-orthophosphate (Perkin Elmer) for 2 h. Medium was then replaced by isotope-free medium and cells were harvested directly in TriPure Reagent (Roche) 3 h post-labelling. Total RNA was extracted using TriPure Reagent standard procedure and dissolved in formamide. 1 µg of total RNA was denatured at 75°C for 10 min and separated in a 1% agarose-formaldehyde-tricine/triethanolamine gel. The gel was dried for 2 h at 80°C under vacuum. Labelled rRNAs were visualised by autoradiography and quantified using a Typhoon PhosphorImager (GE HealthCare). Isotope signal was normalised to ethidium bromide signal for 28S and 18S rRNA bands.

**Preparation of mRNA-associated polysomes**

This was performed as described in [7]. Briefly, cells were seeded at $10^7$ cells/15 cm dish and treated with 5-FU 48 h later as indicated. Cells were then incubated for 5 min with 25 µg/mL emetin (Sigma) and washed twice with cold 1X PBS before harvesting. Cytosolic lysates were prepared as described above. 3 mg of cytosolic proteins was loaded onto a 10-40 % sucrose gradient, sedimented by ultra-centrifugation for 2 h at 240,000 g at 4°C on a SW-41 rotor (Beckman). Fractions were collected and absorbance profiles were generated at 245 nm using an ISCO UA-6 detector. Fractions corresponding to the second half of the polysomes (heaviest polysomes) were pooled and RNAs were extracted with TriPure Reagent as described by the manufacturer (Roche).

**Quantitative RT-PCR**

250 ng of total RNA were reverse transcribed using the M-MLV RT kit and random primers (Invitrogen), according to the manufacturer's instructions. Quantitative real-time PCR (qPCR) was carried out using

the Light cycler 480 II real-time PCR thermocycler (Roche). Expression of mRNAs was quantified using LightCycler 480 SYBR Green I Master Mix (Roche). The primers were obtained by oligonucleotide synthesis (Eurogentec) and are described in Extended data Table 1.

**Xenograft tumour model**

Mice were 7 weeks old female Hsd: Athymic Nude-*Foxn1*$^{nu}$ (Envigo). 1.5×10$^{6}$ HCT116 cells were subcutaneously injected into nude mice flank (n = 5). At day 14, tumours had reached an average of 160 mm$^{3}$. Mice received 3 injections (days 14, 18 and 22) with either vehicle (n = 2) or 5-FU (50 mg/kg, n = 3). 4 h after the last treatment, mice were sacrificed and tumours were collected for subsequent analysis. Total RNA was isolated using the TRI REAGENT protocol (SIGMA T9424) followed by a clean-up step (RNeasy Micro, QIAGEN). rRNA were gel-purified as described above. Animal experiments were performed within French guidelines for experimental animal studies, under DSV agreement A34-172-13.

**Colorectal human tumour samples**

Patient tumours were collected and snap frozen. Total RNA was isolated using the TRI REAGENT protocol (SIGMA T9424) followed by a clean-up step (RNeasy Micro, QIAGEN). rRNAs were gel-purified as described above. Human samples were used under clinical agreement #NCT01577511 under authority of Nîmes Carrémeau University Hospital. All patients signed an informed consent.

**In vitro hybrid translation**

Hybrid *in vitro* translation assay was performed as described previously (35) and is summarised hereafter. After centrifugation of 1 mL of RRL for 2 h 15 min at 240,000 g, 900 µL of ribosome-free RRL (named S100) was collected, frozen and stored at -80 °C. The extent of ribosome depletion from reticulocyte lysate was checked by translating 27 nM of *in vitro* transcribed capped and polyadenylated globin-Renilla mRNA in the S100 RRL and validated when no luciferase activity could be detected. In parallel, transfected cells were lysed in hypotonic buffer R (HEPES 10 mM pH 7.5, $CH_3CO_2K$ 10 mM, $(CH_3CO_2)_2Mg$ 1 mM, DTT 1 mM) and potter homogenised (around 100 strokes). Cytoplasmic fraction was obtained by 13,000 g centrifugation for 10 min at 4°C. The ribosomal pellet was then obtained by ultracentrifugation for 2 h 15 min at 240,000 g in a 1 M sucrose cushion and was rinsed three times in buffer R2 containing HEPES 20 mM, NaCl 10 mM, KCl 25 mM, $MgCl_2$ 1.1 mM, β-mercapto-ethanol 7 mM and resuspended in 30 µL of buffer R2 to reach a ribosomal concentration exceeding 10 µg/µL for optimal and long-term storage at -80°C. The reconstituted lysate was then assembled by mixing 5 µL of S100 RRL with a scale from 0.25 to 4 µg of ribosomal pellet. Typically, the standard reaction contained 5 µL of ribosome-free RRL with 1 µg ribosomal pellet in a final volume of 10 µL. Upon reconstitution, the translation mixture was supplemented with 75 mM KCl, 0.75 mM $MgCl_2$ and 20 µM amino-acid mix.

For *in vitro* translation assays, p0-Renilla vectors containing the β-globin, GAPDH 5'UTR, CrPV, DCV or EMCV IRESs were described previously [38]. mRNAs were obtained by *in vitro* transcription, using 1 µg of DNA templates linearized at the AflII sites, 20 U of T7 RNA polymerase (Promega), 40 U of RNAsin (Promega), 1.6 mM of each ribonucleotide triphosphate, 3 mM DTT in transcription buffer containing 40 mM Tris-HCl (pH 7.9), 6 mM MgCl2, 2 mM spermidine and 10 mM NaCl. For capped mRNAs, the GTP concentration was reduced to 0.32 mM and 1.28 mM of m7GpppG cap analogue (for β-globin mRNA) or m7GpppA (for CrPV mRNA) (New England Biolabs) was added. The transcription reaction was carried out at 37°C for 2 h, the mixture was treated with DNAse and the mRNAs were precipitated with ammonium acetate at a final concentration of 2.5 M. The mRNA pellet was then resuspended in 30 µL of RNAse-free water and mRNA concentration was determined by absorbance using the Nanodrop technology. mRNA integrity was checked by electrophoresis on non-denaturing agarose gel.

**Cell growth and viability assays**

HCT116 were seeded onto 96-well plates at 3,000 cells / well. 48 h after seeding, cells were treated with either 10 µM 5-FU alone or in combination with 5 µM NVP-AEW541 (Sigma-Aldrich), or with DMSO as a control. MTS assay was carried out 48 h after treatment. Cell viability was evaluated using the MTS assay (Cell Titer Aqueous One Solution Cell Proliferation Assay, Promega) according to the manufacturer's protocol. Cell growth was monitored in real time using the xCELLigence technology (ACEA Biosciences), based on electric impedance generated by cells attached to the well. Signals were normalised against the time obtained with IGF-1 (5 or 10 ng/mL (Peprotech)) treatment. Growth rate was calculated over 72 h as the slope under the curve.

**High throughput automated live cell counting by HCS (High Content Screening)**

Cell counting by HCS was performed on a Perkin Elmer Operetta-CLS system, with a 20x NA 0.4 objective. Analysis was performed on 25 fields per well. Hoechst 33342 fluorescence was measured using a 355-385 nm filter for excitation and a 430-500 nm filter for emission. Image analysis and nucleus counting were performed with the Columbus software (Perkin-Elmer).

**Statistical analysis**

Statistical analysis was performed using the Prism software (version 7.0. GraphPad). A two-tailed unpaired student t-test was used for evaluating significance.

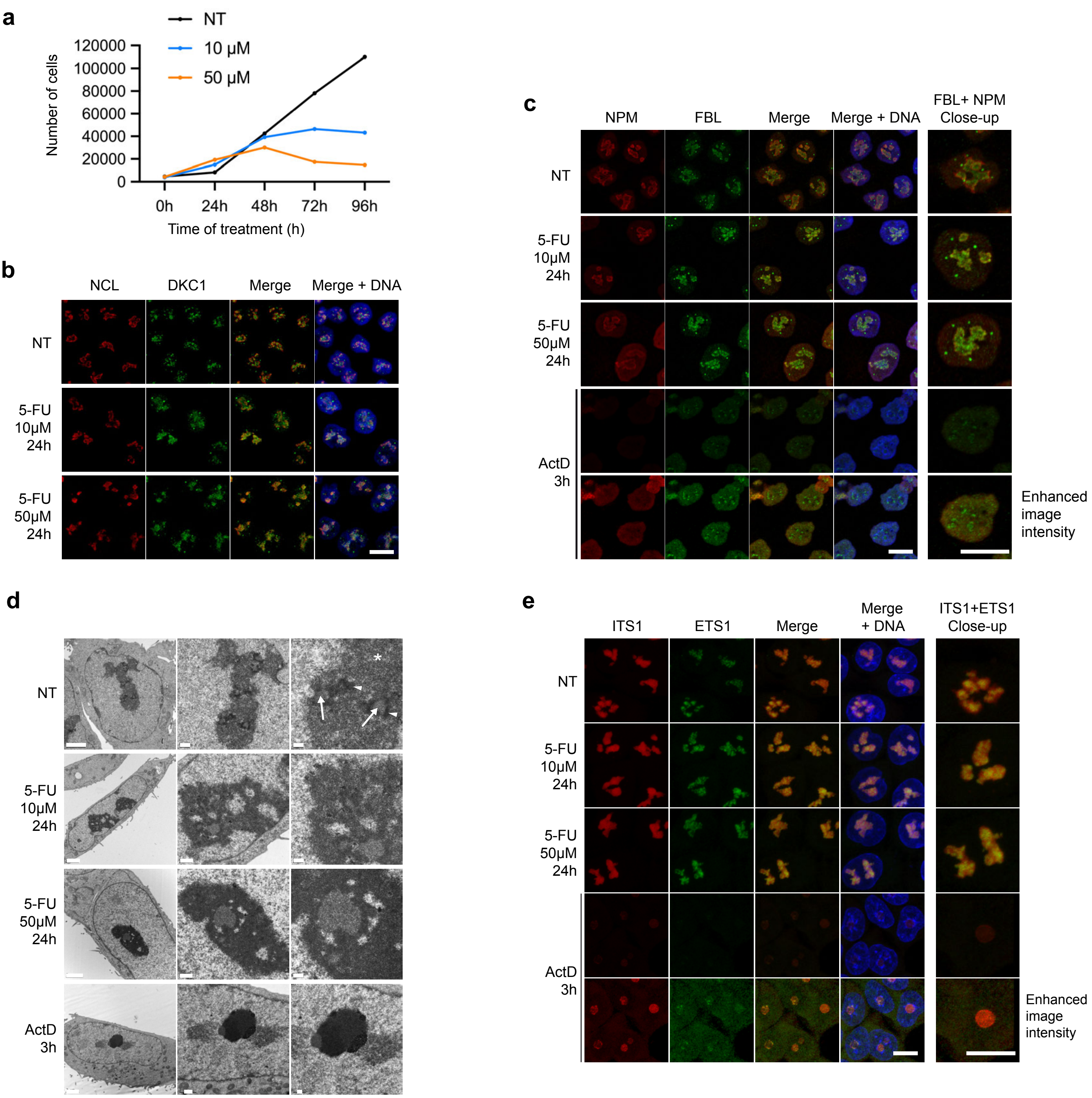


**Extended data Figure 1, related to Figure 1: Ribosome production is maintained in 5-FU treated cells.**
**a,** HCT116 were either untreated (NT, black) or treated with 10 µM (light blue) or 50 µM (orange) of 5-FU for the indicated time. Cells were stained with Hoechst and nuclei were counted by high content analysis imaging. Data are number of cells per well. **b-e,** HCT116 cells were either untreated (NT) or treated with 10 µM 5-FU for 24 h, or with 50 ng/mL actinomycin D for 3 h (ActD). **b,c,** Confocal microscopy imaging following immunofluorescence detection of nucleolin (NCL) fibrillarin (FBL), nucleophosmin (NPM) and dyskerin (DKC1). DNA was stained with Hoechst (blue). The right column shows a close-up of one nucleus. To illustrate the dispersal of NPM upon actinomycin D treatment the signal was enhanced (bottom row). The images show that under 5-FU, nucleoli are enlarged, FBL and DKC1 did not reorganise as nucleolar cap and NCL and NPM did not disperse in the nucleoplasm, as it is the case when pre-rRNA synthesis is inhibited (see ActD as a control). These observations are consistent with a maintenance of pre-rRNA synthesis and alteration of processing. **d**, Nucleolus ultra-structure imaged by transmission electron microscopy. Arrows = Fibrillar center; arrow heads = Dense fibrillar component; star = Granular component. Scale: left column = 2 µm, middle column = 0.5 µm, right column = 0.2 µm. TEM data shows that nucleoli ultra-structure is modified and reflects an alteration of ribosome maturation, and not an inhibition of pre-rRNA synthesis. **e,** Confocal microscopy imaging following fluorescent in situ hybridization (FISH) with probes detecting only the 5'-ETS and ITS1 regions of the pre-rRNAs (see Extended data Fig. 2a). DNA was stained with Hoechst (blue). The right column shows a close-up of one nucleus. The ITS1 probe detect several pre-rRNA species and serves as a reference. The 5'ETS probe detect only the 47S and 45S pre-rRNAs and reflect rRNA synthesis. Inhibition of pre-rRNA synthesis by actinomycin D induce a loss of signal with both 5'-ETS and ITS1 probes. In contrast, upon 5-FU treatment, the signal from both probes was detected at level similar to the untreated control cells. These observations are consistent with a maintenance of pre-rRNA synthesis.

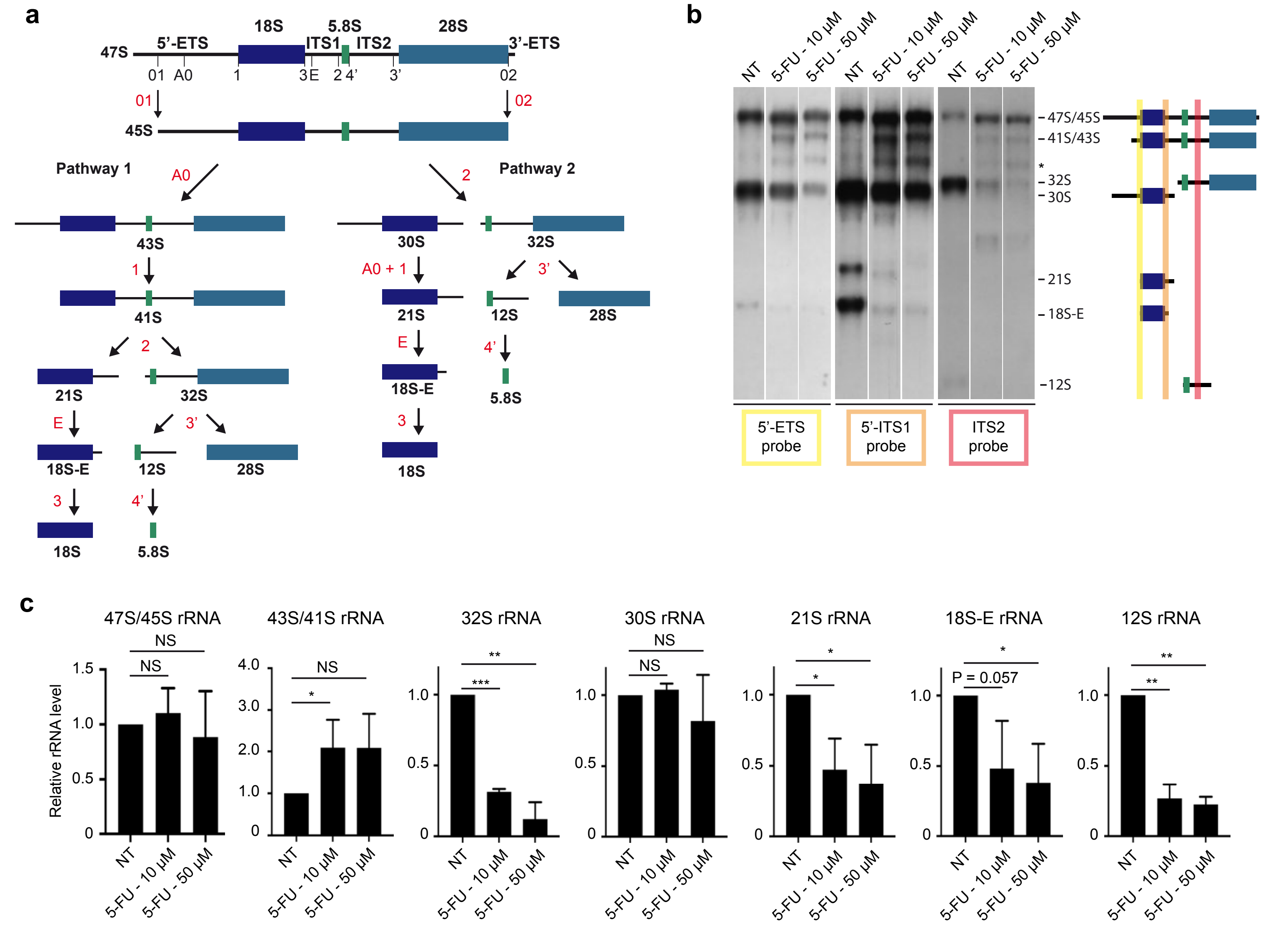


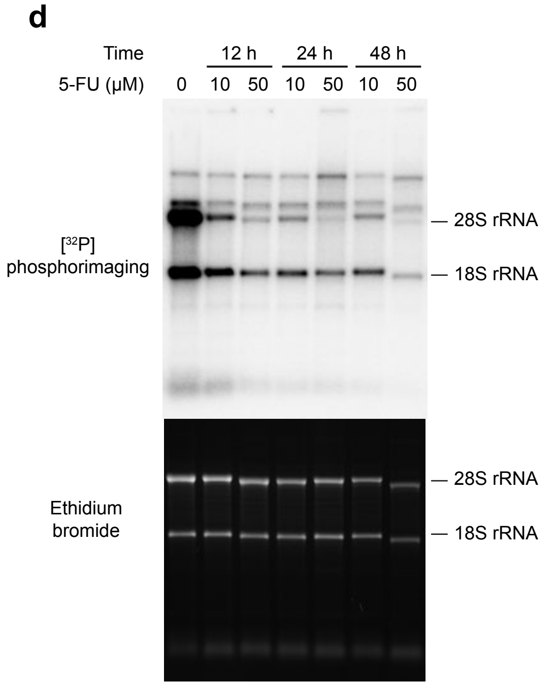


**Extended data Figure 2, related to Figure 1: Ribosome production is maintained in 5-FU treated cells.**

**a**, Simplified representation of pre-rRNA processing adapted from 1. Cleavage sites are indicated in black on the full length 47S pre-rRNA. The products of each cleavage site (in red) are indicated.

**b**, Northern-blot analysis on whole cell RNA from HCT116 cells using probes detecting 5'-ETS, ITS1 or ITS2 spacer regions. The position of the probes and the detected pre-rRNA intermediates are indicated on the right. The star indicates a pre-rRNA intermediate not yet identified.

**c**, Quantification of rRNA intermediates detected by northern-blot, from three independent experiments. Signals were normalized against the intensity of 18S rRNA band detected by ethidium bromide. Data are expressed as mean +/- s.d. n = 3. Results of unpaired two-tailed t-test are indicated by NS = non-significant, * $p<0.05$, ** $p<0.01$, and *** $p<0.001$. These data show that the 47S/45S precursors are produced at similar level in 5-FU treated cells compared to non-treated cells. Increase of 43S/41S intermediates and decrease in 32S and 21S intermediates reflect a processing defect at cleavage site 2.

**d**, Representative images of [$^{32}$P] pulse-chase analysis presented in Fig. 1C. Position of 18S and 28S rRNAs are indicted on the right.

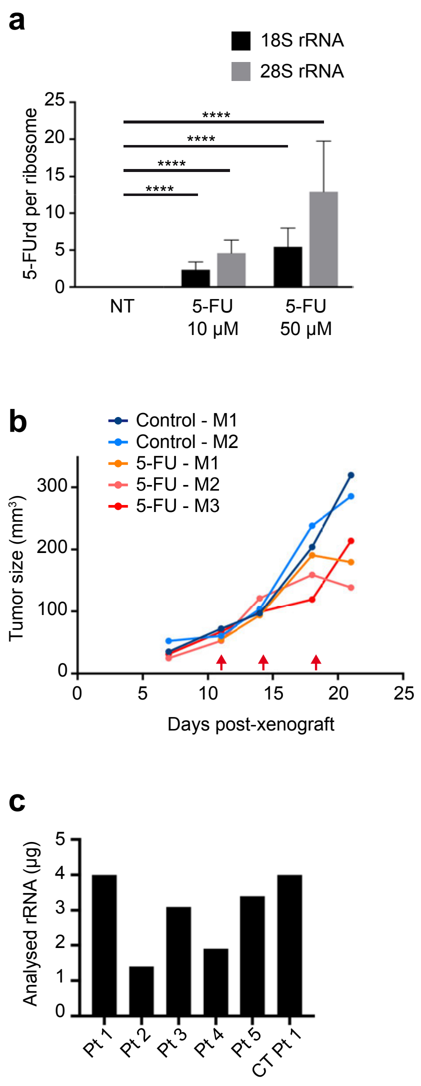


**Extended data Figure 3, related to Figure 2: incorporation of 5-FU in cell lines, xenografts and human tumours.**

**a,** Quantification of 5-FUrd in rRNA by LC-HRMS from 18S and 28S rRNAs that were gel-purified. This approach provides a level of 5-FUrd incorporation similar to that of rRNA extracted from purified ribosomes. Data are from 3 independent experiments. Data are expressed as mean +/- s.d. n = 3. Results of unpaired two-tailed t-test are indicated (**** = p<0.0001).

**b**, HCT116 Xenograft growth curves for individual tumours. Tumor size was measured weekly and are shown in $mm^3$. The results showed that in response to 5-FU treatment, tumour growth is slowed-down.

**c**, Analysis of human colorectal tumour samples. Quantity of rRNA (µg) used in LC-HRMS analysis. For all samples, the quantity of analysed RNA was higher than for our standard protocol (1 µg or rRNA).

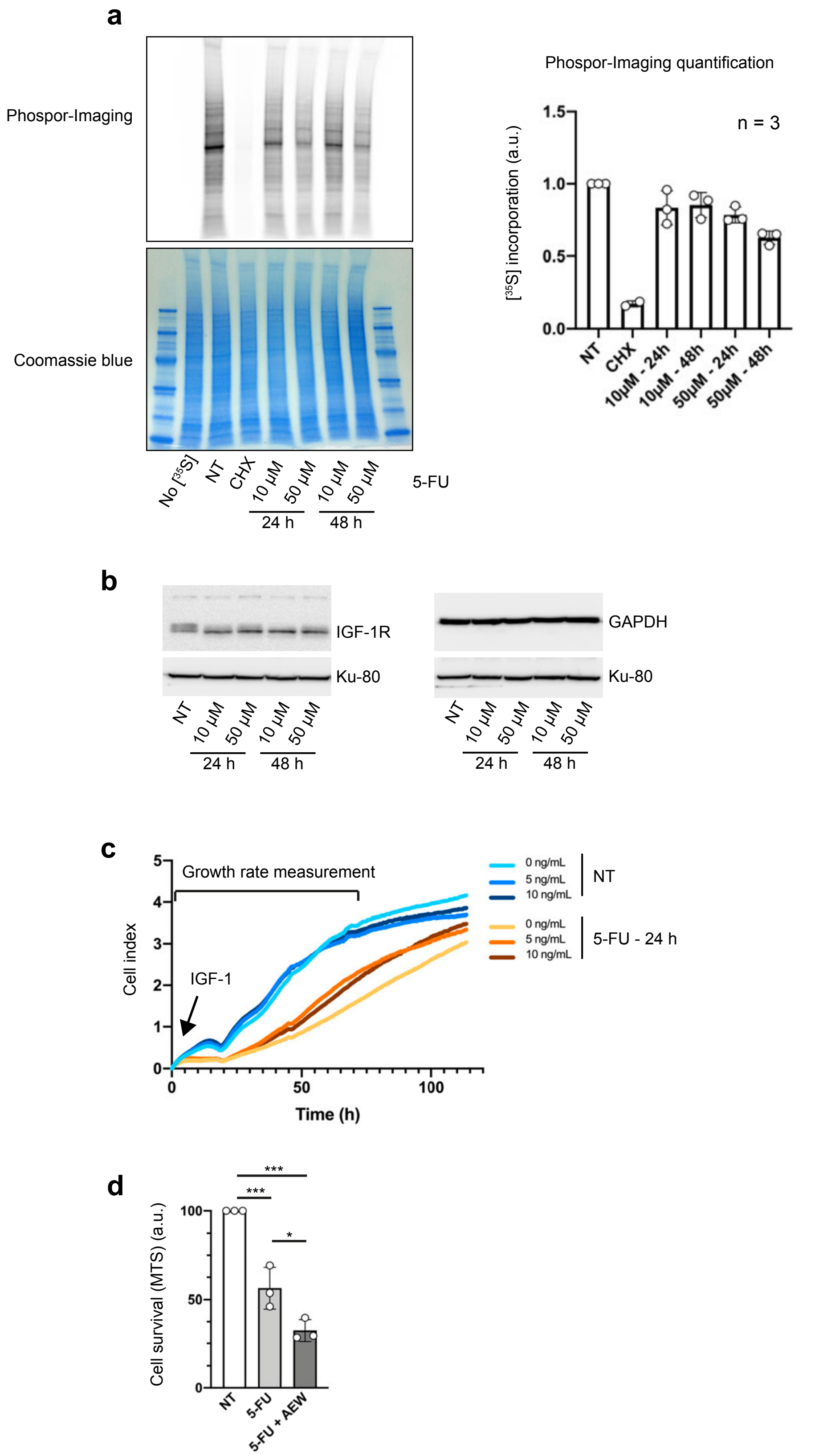


**Extended data Fig. 4. IGF-1R contributes to survival and recovery of 5-FU treated CRC cells.**

**a,** Measure of global protein synthesis by [$^{35}$S]-Met-Cys pulse labelling. HCT116 cells were treated with 10 µM or 50 µM 5-FU for 24 h or 48 h or untreated (NT) or treated with protein synthesis inhibitor cycloheximide at 50 µg/mL for 30 min (CHX). Cells were pulse-labelled with [$^{35}$S]-Met-Cys) for 30 min before cell harvesting, except in the negative control (No [35S]). 10 µg of total protein was analysed by SDS-PAGE (left), and the SDS-PAGE gel was dried and exposed to phosphor-imaging screen (top left). The radioactivity signal was quantified from phosphor-imaging, and was plotted (right). Each dot represents an individual biological sample measured in duplicate and data are expressed as mean ± s.d.

**b,** Same experiment as Fig. 4b. Shown are images of a representative Western blot for the quantification of IGF-1R (left) and GAPDH (right) and using Ku80 as loading control.

**c,** Same experiment as Fig. 4c and d. Shown are individual cell growth curves for each condition. The period used to calculate the growth rate (Fig. 4d) is indicated on top. Each curve is the mean of a technical triplicate (No IGF-1, n = 2 ; IGF-1 treated, n = 3).

**d,** HCT116 cells were treated with 10 µM of 5-FU alone or with 5 µM of IGF-1R inhibitor NVP-AEW541 for 48 h, or untreated (NT). Cell survival was evaluated by MTS assay. Each dot represents a biological replicate and data are mean ± s.d. of a representative experient.

Results of unpaired two-tailed t-test are indicated by * $p<0.05$, ** $p<0.01$, *** $p<0.001$ and **** $p<0.0001$
